# Comparison of musculoskeletal robot biomechanical properties to human participants using motion study

**DOI:** 10.1101/2024.06.27.599434

**Authors:** Iain L. Sander, Aidan C. Sander, Julie A. Stebbins, Andrew J. Carr, Pierre-Alexis Mouthuy

## Abstract

Advanced robotic systems that replicate musculoskeletal structure and function have significant potential for a wide range of applications. Although they are proposed to be better platforms for biomedical applications, little is known about how well current musculoskeletal humanoid systems mimic the motion and force profiles of humans. This is particularly relevant to the field of tendon tissue engineering, where engineered grafts require advanced bioreactor systems that accurately replicate the kinetic and kinematic profiles experienced by the humans *in vivo*. A motion study was conducted comparing the kinetic and kinematic profiles produced by a musculoskeletal humanoid robot shoulder to a group of human participants completing abduction/adduction tasks. Results from the study indicate that the humanoid arm can be programed to either replicate the kinematic profile or the kinetic profile of human participants during task completion, but not both simultaneously. This study supports the use of humanoid robots for applications such as tissue engineering and highlights suggestions to further enhance the physiologic relevance of musculoskeletal humanoid robotic platforms.

## 1. Introduction

Musculoskeletal (MSK) humanoid robots attempt to imitate the body proportion, skeletal structure, anatomical muscle arrangement, and joint function of humans [1]. They have been proposed as a platform for multiple biomedical applications, including exoskeleton training for rehabilitation, production of viable grafts for tendon tissue engineering, and development of advanced orthopedic implants [2-5].The joint performance of MSK robots is of particular interest in developing physiologically relevant models as previous anatomically inspired robots, such as Atlas, rely on rigid joints that do not adequately represent human joint movement or function.

Tendon-driven myorobotic actuation systems are commonly used to mimic muscle action in several MSK humanoid robots, including Roboy, Kenshiro, and Eccerobot [1, 6]. These systems consist of a myorobotic actuator with a brushless dc motor to generate axial tension similar to human muscle fibers, attachment cables which act as the tendon unit, and a motor driver board with a spring encoder to act as the neurological system by sensing relevant variables, including tension, compression, and muscle length [7]. The myorobotic actuators are strategically positioned based on human anatomy and attempt to replicate physiologically relevant behaviors, such as the tendon stretch reflex and adapting to external forces [8].

The ability of actuators to apply physiologically relevant directional stress to tissues is important for tissue tendon engineering applications. *In vivo* studies have demonstrated that healthy tendon tissue deforms nonuniformly [9-13] and mechanobiology research reveals that tendon cells respond distinctively to different types of loading in a directionally dependent manner [14, 15]. Moreover, mechanical testing indicates that tendon tissue will deform differently corresponding to anatomic location [16, 17]. While recent studies have introduced several methods for operating bioreactors, these methods often lack the level of physiological relevance required for effective tissue engineering [3].

MSK robots presented in the literature have been described as replicating the human musculoskeletal system with respect to muscle configuration, anthropometric proportions, joint degrees of freedom and maximum ranges of motion. While these properties are important to replicate human structure, being able to imitate the kinematic and force profile of humans during movement is crucial if MSK robots are to be used in the context of tissue engineering. While some studies have attempted to base their robot’s movement on human kinematic data [18], there is a lack of research comparing the biomechanical profile of these robots to in-vivo human motion. This comparison may be achieved through motion analysis and musculoskeletal modelling.

Motion analysis allows for the quantitative assessment and characterization of kinematics. While there is no widely available method to directly measure *in vivo* muscle forces, computational MSK modelling software can provide reasonable estimates of internal muscle and joint kinetics during functional tasks [19].

We present a study which attempts to address this gap in literature and explore whether MSK humanoid robots adequately imitate the motion and force profile of humans, to support their use in biomedical engineering applications.

## 2. Materials and Methods

### 2.1 Experimental design

The kinematic and kinetic profiles of an MSK humanoid robot over three joint configurations were compared with a reference set of data from forty healthy human participants (Fig. 1). Research ethics approval for the motion study, “Validation of shoulder model through motion tracking study” (reference number: R65527/RE001), was approved by the University of Oxford’s Central University Research Ethics Committee (CUREC).

**Fig. 1.**
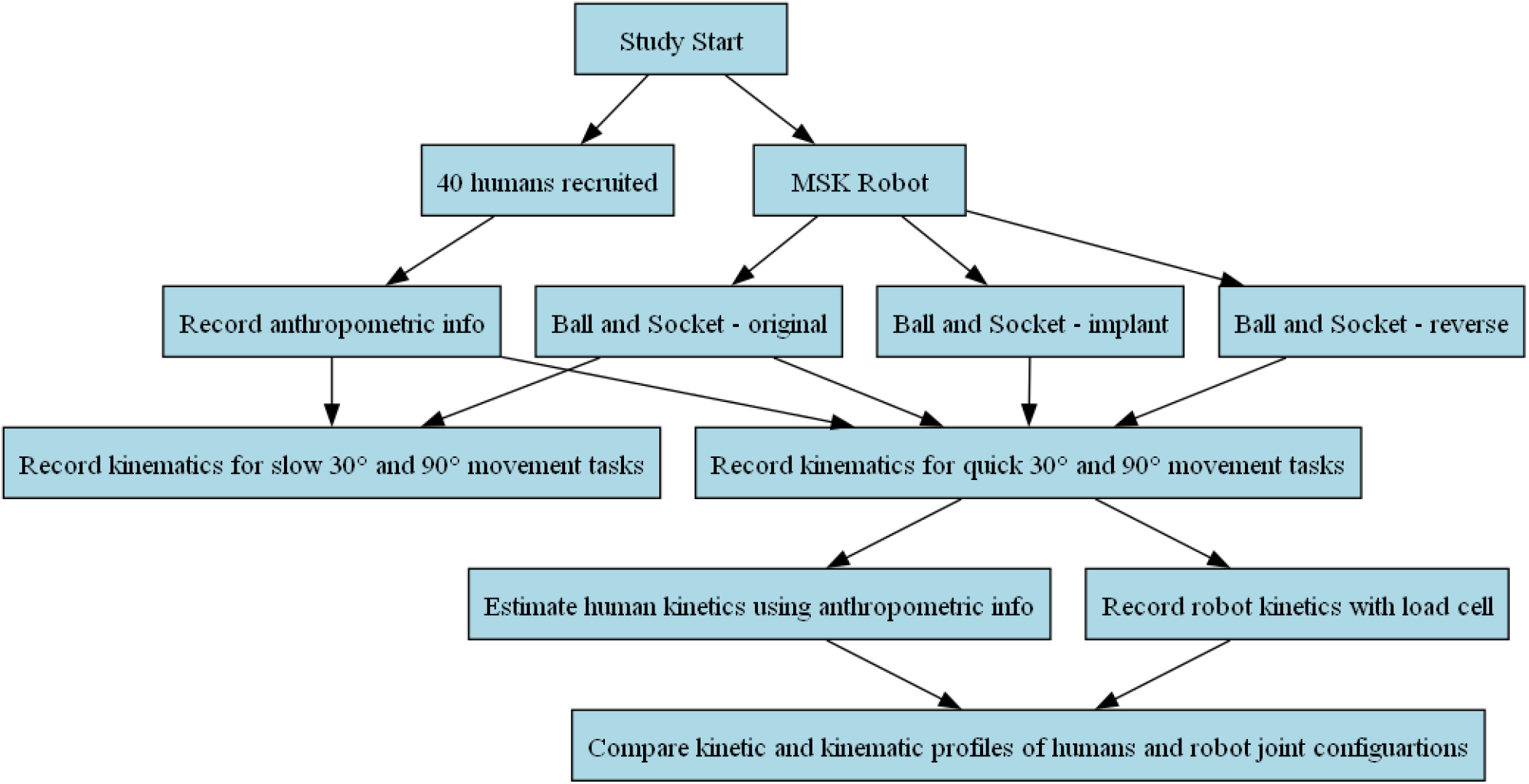
Overview of study procedure and comparison between humans and MSK robot. A group of 40 humans and an MSK robot completed slow (10 deg/s) and fast (30 deg/s) abduction/adduction tasks to 30° and 90°. The MSK robot completed these tasks with three different joint configurations, a ball and socket, clinically used implant, and reverse implant joint. Forces during task completion were estimated in humans using anthropometric information and commercially available modelling software. Forces were directly measured for the humanoid using a load cell. Force was only recorded during the tasks completed at 30 deg/s. Kinematic profiles were captured using Vicon’s Plug-in Gait® for both humans and the MSK robot. All joint configurations completed both tasks to 30° and 90° at the faster speed, with humans and the original ball and socket configuration also completing tasks at the slower speed. Kinetic and kinematic profiles were compared between the humans and humanoid joint configurations.

The study took place at the Oxford Gait Lab in the UK. A Vicon T-Series motion capture system, featuring 16 near-infrared cameras from Vicon Motion Systems (Oxford, UK), was utilized to track retroreflective passive markers (9.5 mm in diameter, B&L Engineering, Santa Ana, USA). Data collection operated at a frequency of 100 Hz, and Vicon NexusTM (v. 2.9.2, Vicon Motion Systems, Oxford, UK) was employed to capture, process, and analyze motion data.

### 2.2 Joint configurations

“Roboy”, the MSK robot whose tendon-driven shoulder component was used throughout this study (Fig. 2A), was developed by Devanthro, GmbH (Garching, Germany). The structural components of the robotic shoulder were constructed using aluminum rods and 3D printed polyamide Nylon 12 (EOS GmbH Electro Optical Systems, Krailling, Germany). The 3D printed structural components were designed using CAD software (Autodesk Fusion, San Francisco, USA). The original joint for this system consisted of a ball component, attached to an aluminum rod to comprise the ‘humerus’ side and a socket component, attached to the main frame of the shoulder, to comprise the ‘thorax’ side of the joint. The original ball and socket joint was developed to resemble the (glenohumeral) GH joint.

**Fig. 2.**
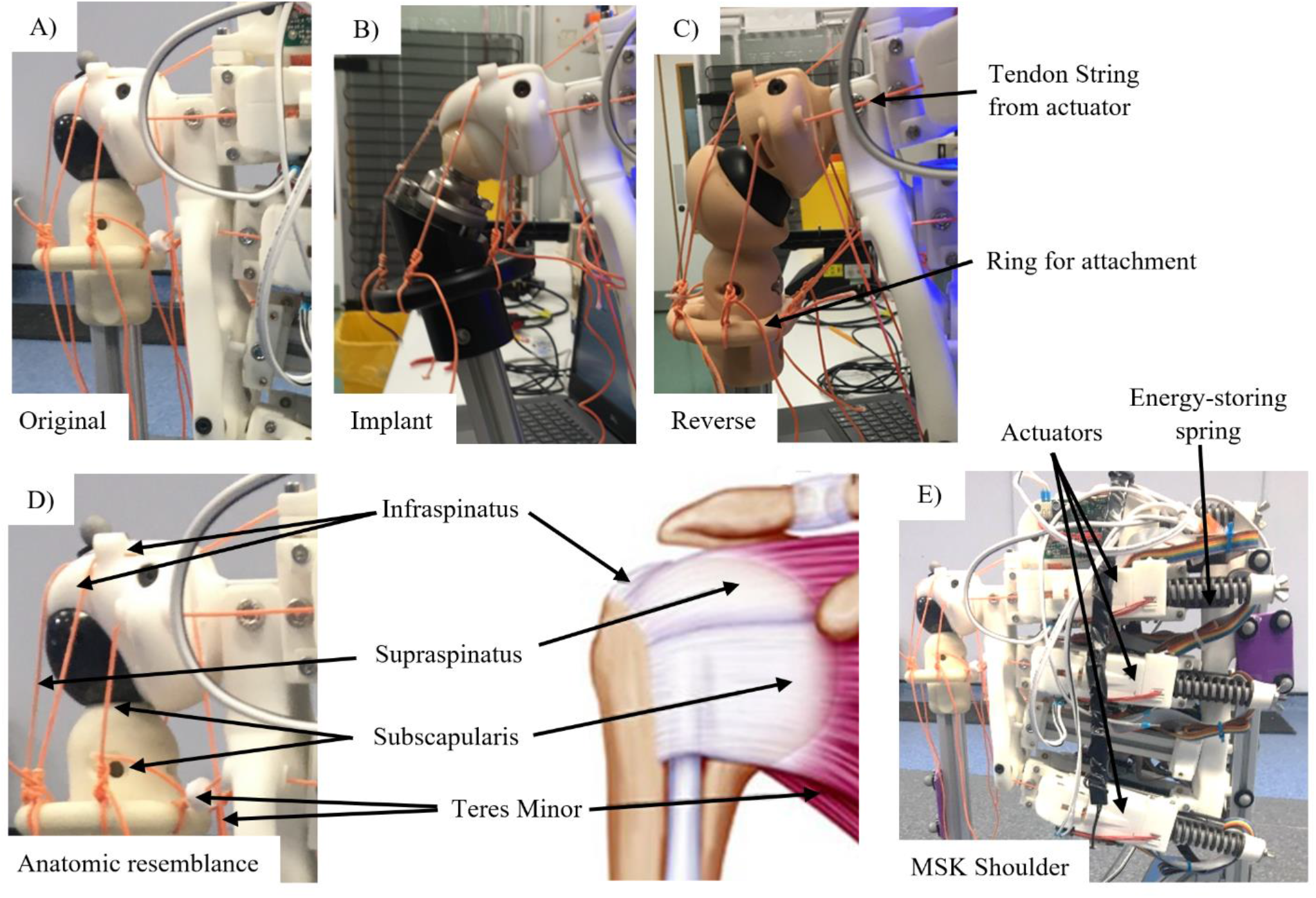
Key components of Roboy arm-shoulder system with ball and socket, reverse shoulder and clinically used implant configuration. A) Original Ball and Socket joint configuration designed by Devanthro. B) Clinically used implant configuration. C) Reverse shoulder configuration. D) Tendon strings are arranged around the joint to imitate the anatomic location of rotator cuff tendons in the human shoulder. The infraspinatus, subscapularis, and teres minor positions on the humanoid arm each have two actuators, for a total of seven actuators contributing to movement. E) Overview of Roboy shoulder and thorax configuration with myorobotic actuators.

Seven myorobotic actuators were arranged to mimic the anatomic location of rotator cuff muscles, controlling joint movements. The subscapularis, teres minor, and infraspinatus muscles were each replicated using two myorobotic actuators. One actuator was used to mimic the supraspinatus. Actuating muscles consisted of a 100 W brushless DC motor (Maxon Motors AG, Sachseln, Switzerland), an encoder, a motor driver board, a spring, and a series of pulleys encased in 3D printed PA 12 [7]. Braided line (300lb Hercules Super Tackle 8 Strands Strong Braid Fishing Line, Hercules, Taiwan) – serving as the tendon – connected the arm of the robotic shoulder to the pulley system and DC motor in each respective actuator. Loading caused the ‘tendon’ line to coil or uncoil around the motor. The movement of this line was discerned by an encoder and interpreted by the motor board driver, with the signal being sent via a serial peripheral interface (SPI) cable to a field programmable gate array (FPGA) board [20]. A proportional-integral-derivative (PID) system was running on the FPGA to control the movement replay speed [7]. ROS Kinetic and Ubuntu 16.04 were the middleware for the FPGA. Rviz, a visualization tool for ROS, was the graphical user interface used to record and replay trajectories.

Two additional configurations were developed to improve the clinical relevance of the original humanoid robot ball and socket joint: a clinically used shoulder implant and a reverse joint inspired by reverse shoulder implants [20]. The purpose of the implant was to evaluate whether the original ball and socket joint reasonably approximated the function of clinical prosthetics. Reverse shoulder arthroplasties are used clinically to improve shoulder function for patients with unstable shoulders and limited abduction due to damaged stabilizing structures around the glenohumeral joint, such as torn rotator cuff muscles or severe osteoarthritis. Given that the humanoid robot lacks a deltoid muscle and stabilizing ligaments around the glenohumeral joint, a reverse shoulder joint was included in this study as it was expected that this configuration would have greater range of motion (ROM) compared to the original ball and socket joint.

### 2.3 Kinematic modeling

Our retroreflective marker protocol for the kinematic modelling was an adaptation of the upper body Plugin Gait® (PiG) model by Vicon [21]. This protocol comprises three segments (thorax, humerus, and forearm) and two joints (shoulder with 3 degrees of freedom and elbow with 2 degrees of freedom). Similar to the upper body PiG, we approximated shoulder kinematics to the glenohumeral (GH) joint. The GH joint was the shoulder joint center in the model, excluding scapular contributions to motion. PiG’s default segment axes and angle calculation methods were employed.

In contrast to the PiG model, our marker protocol featured a cluster on the right upper arm and one under the clavicle marker on the torso. Notably, markers for the head, hands, and left arm were not utilized in this study. For the humanoid robot, markers for the wrist, back, and left shoulder were omitted, given that the system only included a right shoulder joint. Refer to Fig. 3 for specific marker locations and descriptions.

**Fig. 3.**
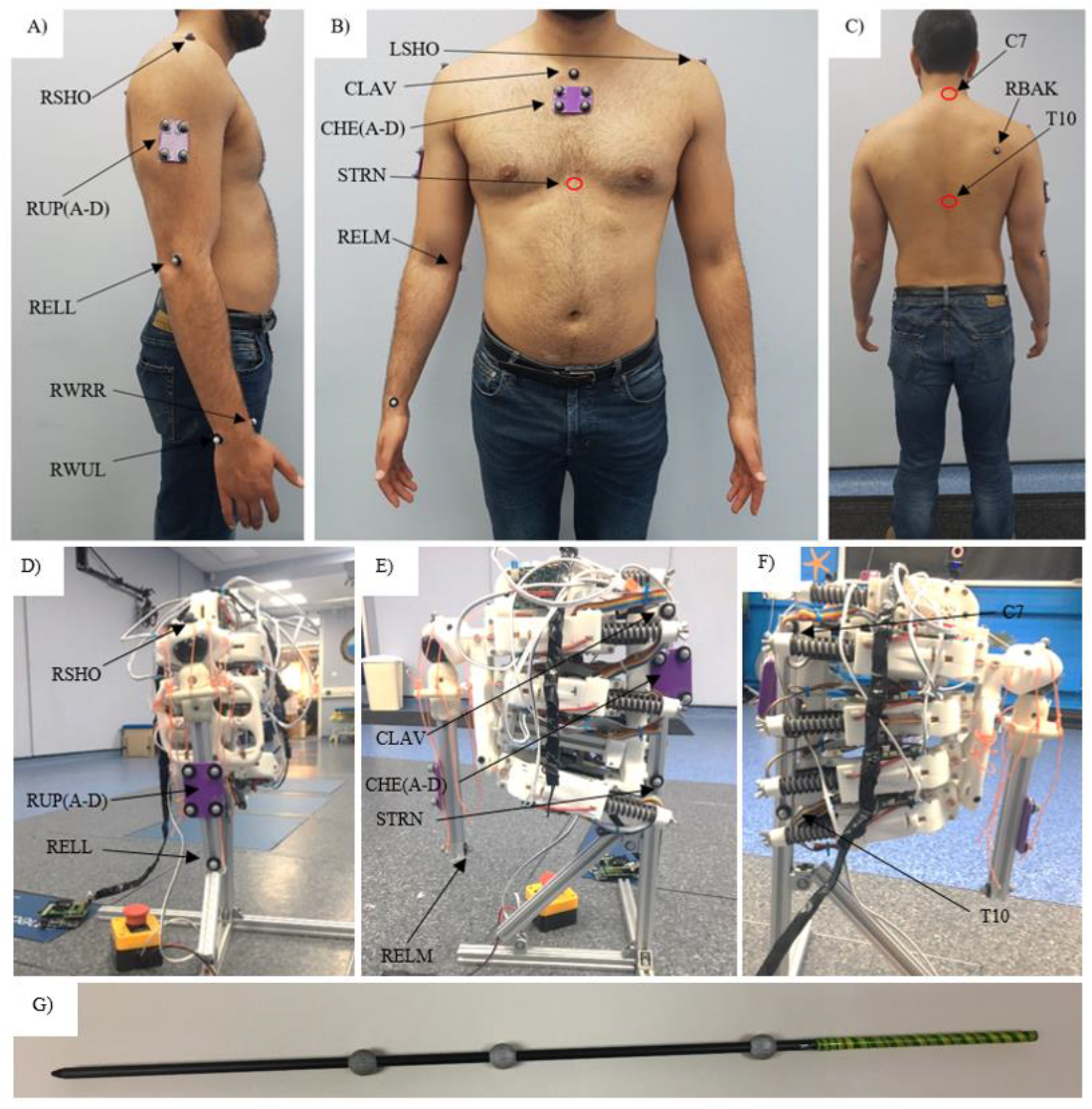
Marker placement protocol used in study. (A– C) Marker placement locations for human participants. Locations indicated by red ellipses (i.e., STRN, C7, and T10) have markers constructed virtually using a pointer wand. (D – F) Marker placement locations on Roboy arm-shoulder system. LFSO: Left Shoulder, RHSO: Right Shoulder, CLAV: Clavicle, CHE(A-D): Chest Cluster, RUP(A-D): Right Upper Arm Cluster, RELL: Right Elbow (lateral), REM: Right Elbow (medial), RWUL: Right Wrist Marker (ulnar side), RWRR: Right Wrist Marker (radial side), RBAK: Right Back (scapula), T10: 10th Thoracic Vertebra, C7: 7th Cervical Vertebra, STRN: Sternum. (G) Wand used for virtual construction of C7, T10, and STRN markers for human participants.

For human participants, markers were placed on anatomical landmarks detailed in Fig. 3, digitized values for the sternum and C7/T10 vertebrae were estimated using a marker wand (Fig. 2G). Elbow and wrist joint centers were computed as the midpoint between their medial and lateral markers. The shoulder joint center was determined using the method outlined by Rab et al [22].

For the robotic arm-shoulder system, markers were positioned corresponding to human anatomical landmarks. The robot’s elbow joint center was determined as the midpoint between the medial and lateral ‘elbow’ markers. As the right shoulder marker for the robotic arm was placed directly superior to the shoulder joint center, the offset from the ‘RHSO’ marker (Fig. 3D) was measured using a clinical caliper. This distance was measured for each joint configuration. The program code accounted for different offsets in shoulder joint centers.

### 2.4 Kinetic modeling

AnyMoCap (v.7.3.0, AnyBody Technology, Aalborg, Denmark) was used to estimate supraspinatus forces for human participants using the AMS (AnyBody Modeling System). Our upper body models included the Delft Shoulder and Elbow Model DSEM) and a rigid body spine model developed by Zee et al. [23, 24].

The AMS default upper extremity (UE) segment inertial parameters were derived from anthropometric data published by Winter [25]. Muscles were represented using a simplified Hill-type muscle model, complemented by a 3rd order polynomial recruitment and load-sharing algorithm [26]. Within the AMS, muscles were divided into smaller segments, and total muscle forces were computed as the sum of all segments for a specific muscle.

The AnyMoCap application utilized linear model scaling for kinetic estimation [27, 28]. Participant-specific height and weight data were directly inputted into the AMS, and imported data noise was filtered using a second-order zero-phase Butterworth filter with a 10 Hz cut-off frequency.

An optimization process within the AMS, utilizing the filtered motion study data and participant anthropometric information, rescaled the UE model and recalculated joint angles. Inverse dynamics analysis was conducted to estimate joint moments, reaction forces, and muscle forces. The force estimates were specifically tailored for movement tasks at 30 degs/s, aligning with the abduction exercises performed by physical therapists during the later stages of shoulder rehabilitation [29-33].

These force measurements for the humanoid robot were captured using a load cell (model WMC-450N, Interface Force Measurements Ltd, Crowthorne, UK) and documented with a high-speed data logger (model 9330-MSDI-IP43, Interface Force Measurements Ltd, Crowthorne, UK) at a frequency of 10 Hz. Force data for the humanoid robot was only recorded for the 30 deg/s movement trials. The load cell was affixed to the robotic system on the ‘humeral’ side and connected to an actuating muscle in the ‘supraspinatus’ position using braided ‘tendon’ line (300lb Hercules Super Tackle 8 Strands Strong Braid Fishing Line, Hercules, Taiwan).

### 2.5 Participants

Forty participants (23 male, 17 female; mean age: 28.3 years, range: 20.5 – 36.9 years; mean height: 1.75 m, range: 1.56 – 1.93 m) were recruited from the University of Oxford, volunteering for the study without compensation. Inclusion criteria required participants to be older than 18 and have no neurological or musculoskeletal pathologies that would limit task completion. Candidates with a history of upper extremity surgery, shoulder injuries within the past year, or any physical disabilities or injuries impeding shoulder movement were excluded from the study.

Participants completed a measurement form which recorded their age, gender, height, and weight, measured using a stadiometer and a scale (EKS Scales, Hong Kong). Participant IDs were pseudonymized. One participant’s data was excluded from analysis due to poor data quality.

### 2.6 Task protocol

Tasks for this study were completed with the right arm only. The supraspinatus is primarily responsible for abduction up to 15° and assists the deltoid with motion up to 90° in healthy shoulders [34]. The supraspinatus is most active during the initial phases of abduction from 0° to 30°, contributing significantly to abduction up until 90° [35]. These ranges are clinically significant because they represent the ROM most frequently involved in daily activities and shoulder injury rehabilitation exercises [29-33].

Tasks involved two movement speeds: a slow, controlled motion over 6 seconds for the 30° task and 18 seconds for the 90° task (targeting 10 deg/s), as well as a faster motion over 2 seconds for the 30° task and 6 seconds for the 90° task (targeting 30 deg/s). These timings allowed for the identification of limitations in the humanoid arm’s movement speed and its ability to mimic early and late-stage rehabilitation movements. Slow, controlled motions typically align with early phases of physical therapy following injury, whereas the faster isokinetic movements mimic healthy shoulder function [36, 37].

One researcher conducted data collection for both human and robotic trials to maintain marker placement consistency. Human participants received standardized instructions and a demonstration. The assessor used an on-screen stopwatch and verbal counting to ensure timing reliability. Each participant had one representative trial recorded per task, aligning with prior literature recommending a single representative trial for clinical decision-making in motion analysis studies [38, 39].

For the robotic arm trials, the assessor recorded a movement to the desired range and speed, programming the robotic arm to replicate it. This recording involved manual arm manipulation to achieve the desired range and angular velocity. The robotic arm control software was configured to reproduce these movements.

Each task was programmed five times at 10 deg/s and twenty times at 30 deg/s for the original ball and socket joint. The clinically used shoulder implant and reverse ball and socket joint only completed the “abduction to 30 degrees” and “abduction to 90 degrees” tasks at 30 deg/s. The same PID controller settings were applied to the robotic system for each joint configuration, ensuring consistent performance comparison during task completion.

### 2.7 Kinematic and kinetic profile comparison

Motion trajectories were plotted for all joint configurations. Total range of motion and average movement speed were used to evaluate ability to complete movement tasks. Movement quality was assessed using peak acceleration, a measure of efficiency during task completion, and the speed metric, a dimensionless ratio of average to peak movement speed (v_avg_/v_peak_) [40-44]. A speed metric ratio closer to 1 indicates greater consistency of speed throughout a motion, while a value closer to 0 indicates larger fluctuations in speed consistency throughout a movement [41, 43].

Kinetic data were presented as force (N) in this study to facilitate comparison between human participants and the humanoid robot. Force trajectory plots were presented for qualitative comparison of humanoid robot and participant kinetic profiles. Average force and peak force were the outcome parameters used to quantitatively compare the kinetic profiles of the humanoid arm and healthy participants.

### 2.8 Statistical Analysis

The D’Agostino-Pearson test was used to verify the normality of outcome parameters [45]. Kinematic outcome parameters were found to be normally distributed for both the humanoid robot and healthy participants. Unpaired t-tests were therefore used for quantitative analysis of kinematic data. Force outcome parameters obtained from human participants using MSK modelling were normally distributed; several force values obtained from the humanoid robot measured using a load cell were not normally distributed. The Mann-Whitney U was therefore used for quantitative interpretation of kinetic data.

## 3. Results

### 3.1 Comparing humanoid and human movement profiles

The robot’s mean trajectory plot for the 30° abduction/adduction “slow” task is nearly identical to that of humans with the original ball and socket joint (Fig. 3A). There is considerable variability in the motion data for this task, as noted by the wide error band. In the 90° trial, the original joint configuration achieved a slightly smaller total ROM compared to healthy human participants (Fig. 3B).

In the 30° abduction/adduction task, only the robot’s average and peak accelerations are statistically different from humans, being 3.7 (SEM 1.4) deg/s^2^ and 23.1 (SEM 2.6) deg/s^2^ lower, respectively (Table 1). In the 90° task, the robot matched human speed but achieved a 9.4° (SEM 3.1°) smaller range of motion. The average and peak accelerations of the robot are also lower than those of the humans, with differences of 1.8 (SEM 0.7) deg/s^2^ and 15.8 (SEM 3.1) deg/s^2^, respectively.

**Table 1:** Kinematic outcome variables from slow movement 30° and 90° abduction/adduction task. P-values indicating that a parameter of interest differs significantly from human participants (p<0.05) have been bolded.

| Parameter | Human |  | Humanoid Robot<br>(original joint) |  | p-value |  | Difference (SEM) |  |
| --- | --- | --- | --- | --- | --- | --- | --- | --- |
|  | Mean (SD) |  | Mean (SD) |  |  |  |  |  |
|  | 30° | 90° | 30° | 90° | 30° | 90° | 30° | 90° |
| **Total ROM (deg)** | 30.3 (7.0) | 78.1 (6.6) | 30.7 (0.4) | 68.7 (4.3) | 0.702 | **0.0042** | 0.4 (1.1) | 9.4 (3.1) |
| **Average Speed (deg/s)** | 9.7 (2.3) | 7.9 (0.9) | 8.6 (0.3) | 7.5 (0.3) | 0.297 | 0.363 | 1.1 (1.0) | 0.4 (0.4) |
| **Peak Speed (deg/s)** | 16.1 (5.4) | 16.4 (5.3) | 16.1 (0.8) | 13.5 (2.4) | 0.976 | 0.246 | 0.0 (0.9) | 2.8 (2.4) |
| **Speed Metric** | 0.61 (0.1) | 0.52 (0.1) | 0.53 (0.1) | 0.57 (0.1) | 0.190 | 0.346 | 0.1 (0.1) | 0.1 (0.1) |
| **Average Acceleration (deg/s<sup>2</sup>)** | 9.0 (3.2) | 5.7 (1.5) | 5.4 (0.5) | 3.9 (0.7) | **0.0145** | **0.011** | 3.7 (1.4) | 1.8 (0.7) |
| **Peak Acceleration (deg/s<sup>2</sup>)** | 36.0 (16.0) | 33.0 (17.14) | 12.9 (1.1) | 17.2 (2.9) | **<0.0001** | **<0.0001** | 23.1 (2.6) | 15.8 (3.1) |

Fig. 5 shows trajectories for the humanoid robot and human participants performing the abduction/adduction tasks at a faster speed. The peak ROM for the original ball and socket joint (Fig. 4C, F) and implant joint (Fig. 4D,G) were significantly below the target ROM of 30° and 90°. In contrast, the reverse shoulder joint closely matched the target peak ROM of 30° and 90° (Fig. 4 E, H). Additionally, the robot consistently reached its highest ROM before midcycle, while human participants achieved peak abduction range at the midpoint of the cycle (Fig. 5A, B).

**Fig. 4.**
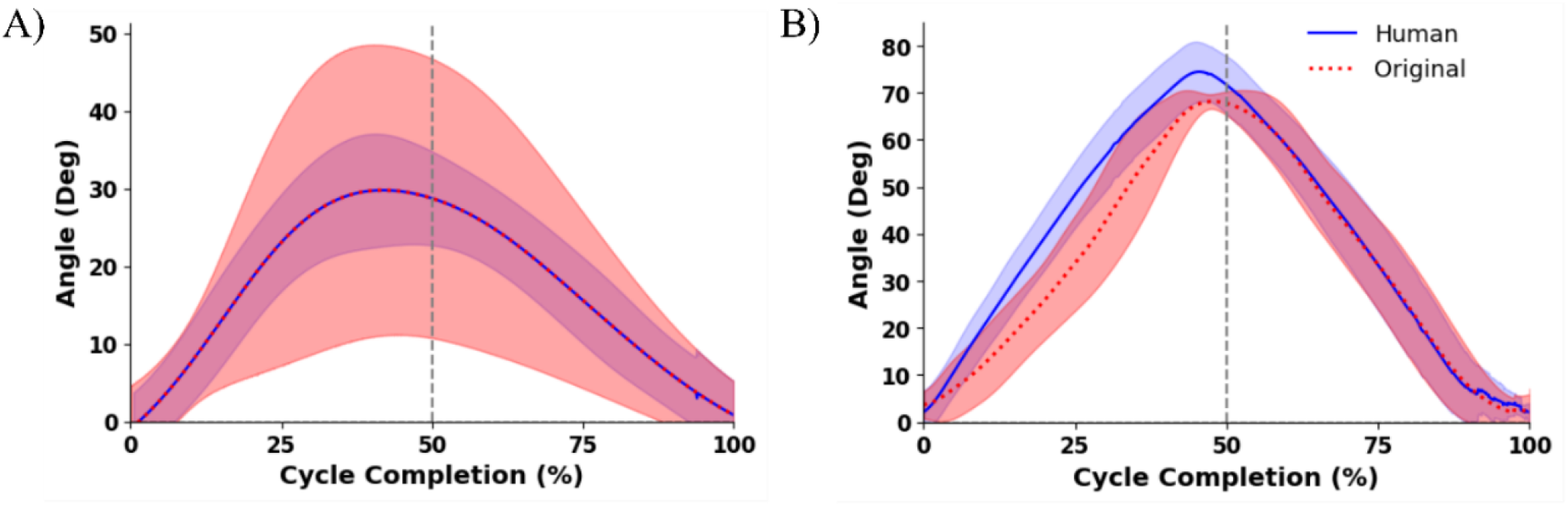
Kinematic trajectory plots for slow movement tasks using the original ball and socket joint configuration. (A) Compares the humanoid arm to human participants for the 30° abduction task. The cycle length of time for this task was 6 s. (B) Compares the humanoid arm to human participants for the 90° abduction task. The cycle length of time for this task was 18 s. The blue curve and error bands represent mean human participant movement ± 1 SD in both plots. The red curve and error bands represent mean angle ± 1 SD for the humanoid robot in both plots.

**Fig. 5.**
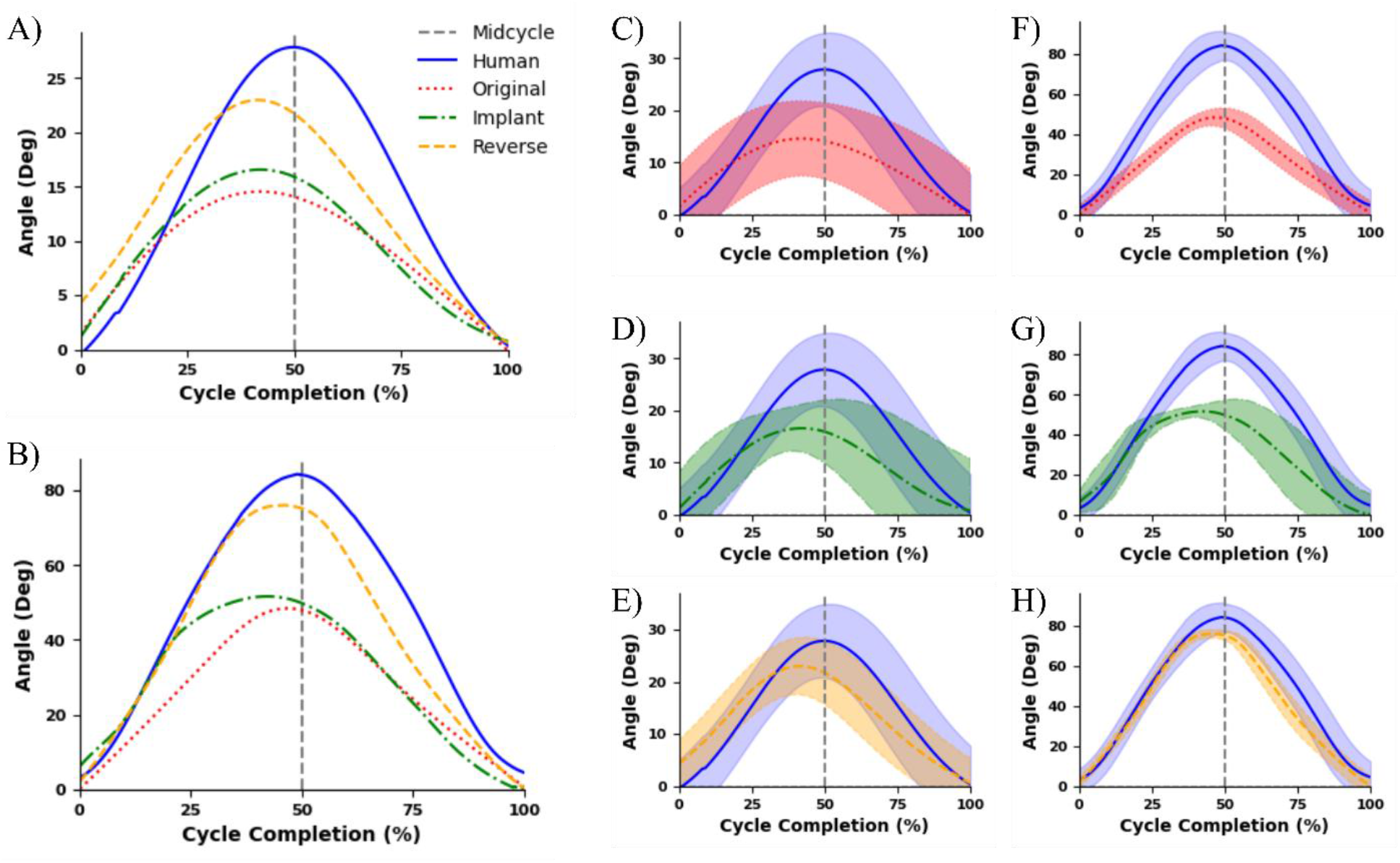
Kinematic trajectory plots for movement to 30° and 90° in the scapular plane, measured in degrees. (A) Compares the trajectory of the original joint configuration, the clinical implant, and reverse shoulder joint configuration over the 2 second 30° movement length without error bands. (C, D, E) Compare the trajectory of the original joint configuration, the clinical implant, and reverse shoulder joint configuration over the 2 second 30° movement length. (The movement cycle length is 2 seconds. (F, G, H) Present the same comparisons with a 90° movement length over 6 seconds. (B) Comparison of the three joint configurations with humans over the 90° movement length without error bands. Blue curves and error bands represent mean human participant movement ± 1 SD. The colored error bands represent mean angle ± 1 SD for each joint configuration.

The clinical implant joint closely matched human speed until reaching the peak abduction range, moving more slowly during the adduction phase (Fig. 5D, G) On the other hand, the original joint was slower compared to humans throughout both tasks (Fig. 5C, F). Qualitatively, the reverse shoulder joint configuration most closely resembled the human speed profiles during both tasks (Fig. 5E, H).

Table 2 summarizes the kinematic outcome parameters for the humanoid robot with different joint configurations during the fast speed task. The original and clinical implant achieved peak ROMs approximately 10° and 30° lower than the target ROMs of human participants for the 30° and 90° abduction tasks. In contrast, the reverse shoulder joint achieved a total ROM within 10° of human participants for both tasks.

**Table 2:** Kinematic outcome variables from 30° and 90° abduction/adduction task used to evaluate implant performance. P-values indicating that a parameter of interest differs significantly from human participants (p<0.05) have been indicated in bolded text.

| **Kinematic outcome variables from 30° abduction/adduction task** |  |  |  |  |  |  |  |  |  |  |
| --- | --- | --- | --- | --- | --- | --- | --- | --- | --- | --- |
| **Parameter** | Human | Robotic shoulder - Original |  |  | Robotic shoulder - Implant |  |  | Robotic shoulder - Reverse |  |  |
|  | Mean (SD) | Mean (SD) | p-value | Difference (SEM) | Mean (SD) | p-value | Difference (SEM) | Mean (SD) | p-value | Difference (SEM) |
| **Total ROM** | 32.5 (6.9) | 16.0 (6.6) | **<0.0001** | 16.5 (1.7) | 21.7 (2.2) | **<0.0001** | 10.8 (1.2) | 24.2 (3.4) | **0.0016** | 8.3 (1.3) |
| **Average Speed (deg/s)** | 29.7 (6.9) | 9.7 (3.2) | **<0.0001** | 20.0 (1.3) | 14.0 (2.5) | **<0.0001** | 15.8 (1.2) | 21.2 (2.8) | **<0.0001** | 8.5 (1.3) |
| **Peak Speed (deg/s)** | 49.7 (14.5) | 19.0 (3.8) | **<0.0001** | 30.7 (2.5) | 30.1 (8.1) | **<0.0001** | 19.6 (3.5) | 34.6 (4.2) | **<0.0001** | 15.1 (2.5) |
| **Speed Metric** | 0.60 (0.11) | 0.51 (0.17) | **0.016** | 0.09 (0.03) | 0.48 (0.1) | **0.0002** | 0.11 (0.03) | 0.63 (0.12) | 0.352 | 0.03 (0.03) |
| **Average Acceleration (deg/s<sup>2</sup>)** | 73.7 (31.0) | 12.0 (2.5) | **<0.0001** | 61.7 (5.0) | 29.4 (10.6) | **<0.0001** | 44.3 (5.5) | 41.5 (9.3) | **<0.0001** | 32.2 (5.4) |
| **Peak Acceleration (deg/s<sup>2</sup>)** | 173.3 (76.2) | 27.0 (7.5) | **<0.0001** | 146.2 (12.3) | 57.8 (20.7) | **<0.0001** | 115.4 (17.4) | 108.1 (34.1) | **<0.0001** | 65.1 (14.4) |
| **Kinematic outcome variables from 90° abduction/adduction task** |  |  |  |  |  |  |  |  |  |  |
| **Total ROM** | 82.6 (8.6) | 50.9 (3.1) | **<0.0001** | 31.7 (2.0) | 54.1 (7.5) | **<0.0001** | 28.5 (2.2) | 77.0 (2.1) | **0.0062** | 5.6 (2.0) |
| **Average Speed (deg/s)** | 26.6 (4.4) | 13.2 (0.7) | **<0.0001** | 13.4 (0.7) | 16.7 (2.1) | **<0.0001** | 9.9 (1.0) | 25.1 (0.7) | 0.130 | 1.5 (1.0) |
| **Peak Speed (deg/s)** | 41.2 (6.7) | 17.6 (1.3) | **<0.0001** | 24.1 (1.5) | 65.4 (16.9) | **<0.0001** | 23.8 (3.9) | 45.8 (18.8) | 0.195 | 4.2 (3.2) |
| **Speed Metric** | 0.62 (0.14) | 0.75 (0.05) | **<0.0001** | 0.13 (0.02) | 0.28 (0.10) | **<0.0001** | 0.34 (0.04) | 0.58 (0.10) | 0.300 | 0.04 (0.04) |
| **Average Acceleration (deg/s<sup>2</sup>)** | 25.8 (6.4) | 5.6 (0.8) | **<0.0001** | 20.2 (1.5) | 30.3 (10.1) | 0.080 | 4.5 (2.5) | 24.8 (22.1) | 0.846 | 1.0 (5.0) |
| **Peak Acceleration (deg/s<sup>2</sup>)** | 102.8 (55.0) | 22.5 (4.6) | **<0.0001** | 80.3 (12.4) | 199.8 (29.4) | **0.0002** | 97.0 (24.3) | 80.7 (20.0) | 0.096 | 22.1 (13.1) |

The average movement speeds for the original and implant joints were significantly lower than those of human participants for both tasks (Table 2). In the 30° abduction trial, there were differences of 20.0 (SEM 1.3) deg/s for the original and 15.8 (SEM 1.2) deg/s for the clinical implant. In the 90° trial, the differences were 13.4 (SEM 0.7) deg/s and 9.9 (SEM 1.0) deg/s, respectively. The mean peak movement speeds for the original joint were less than half the mean peak movement speeds of human participants in both trials. In contrast, the reverse shoulder joint more closely matched the average movement speed of healthy human participants, with no statistically significant difference in the average movement speed for the 90° task and an 8.5° (SD 1.3)° difference for the 30° task.

There is no statistically significant difference between the speed metric of the reverse shoulder joint and that of human participants for both tasks. The implant joint achieved lower average speed metrics compared to human participants for both tasks. The original joint had a lower speed metric for the 30° task and a higher speed metric value for the 90° task when compared to human participants.

In the 30° task, human participants had higher mean peak acceleration than any of the joint configurations tested. The reverse shoulder joint’s peak acceleration was statistically similar to human participants in the 90° task. However, the clinical implant joint had a peak acceleration 97.0 (SEM 24.3) deg/s^2^ greater than human participants. The original joint had a peak acceleration 80.3 (SEM 12.4) deg/s^2^ lower than human participants in the 90° task.

### 3.2 Comparing humanoid and human force profiles

Supraspinatus forces during the faster (2s) 30° abduction task are shown in Fig. 6 and Fig. 7. No kinetic trials were conducted at the slower speed. In the 30° abduction task, force peaks mid-cycle for humans during the 30° abduction task (Fig. 6A-C). The original joint configuration followed a similar pattern (Fig. 6A). In contrast, the implant and reverse shoulder joint (Fig. 6 C, B) reach peak force levels at the quarter-cycle for the 30° task. In both the 30° abduction tasks, the implant and original joint configurations closely match the force magnitudes of human participants, while the reverse shoulder comparatively generates significantly higher supraspinatus muscle forces throughout the movement cycle.

**Fig. 6.**
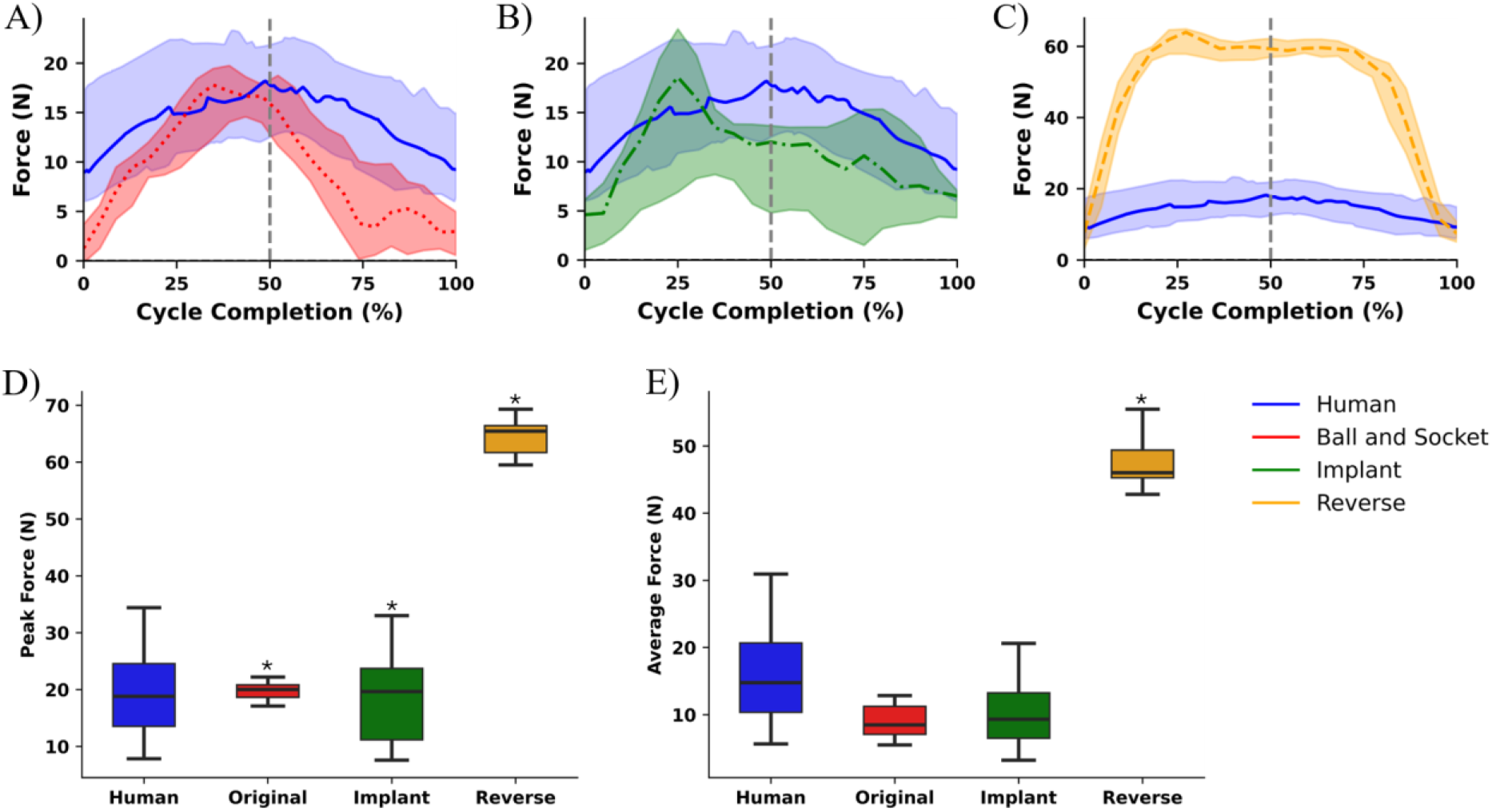
Estimated supraspinatus force profile and values for abduction to 30° in scapular plane, measured in Newtons (N). (A) Compares the original ball and socket joint configuration’s force profile to the human shoulder. (B) Compares the reverse shoulder joint configuration and human shoulder force profiles. (C) Compares the clinical implant joint configuration and human shoulder force profiles. The colored error bands represent the IQR for each joint configuration. (D) Average force values for the human participants and the humanoid robot for each joint configuration during task completion. (E) Peak force values for human participants and the humanoid robot for each joint configuration during task completion. Results which have statistically significant (p<0.05) values from human participants have been denoted with an asterisk (*). The movement cycle length is 2 seconds corresponding the “fast” condition.

**Fig. 7.**
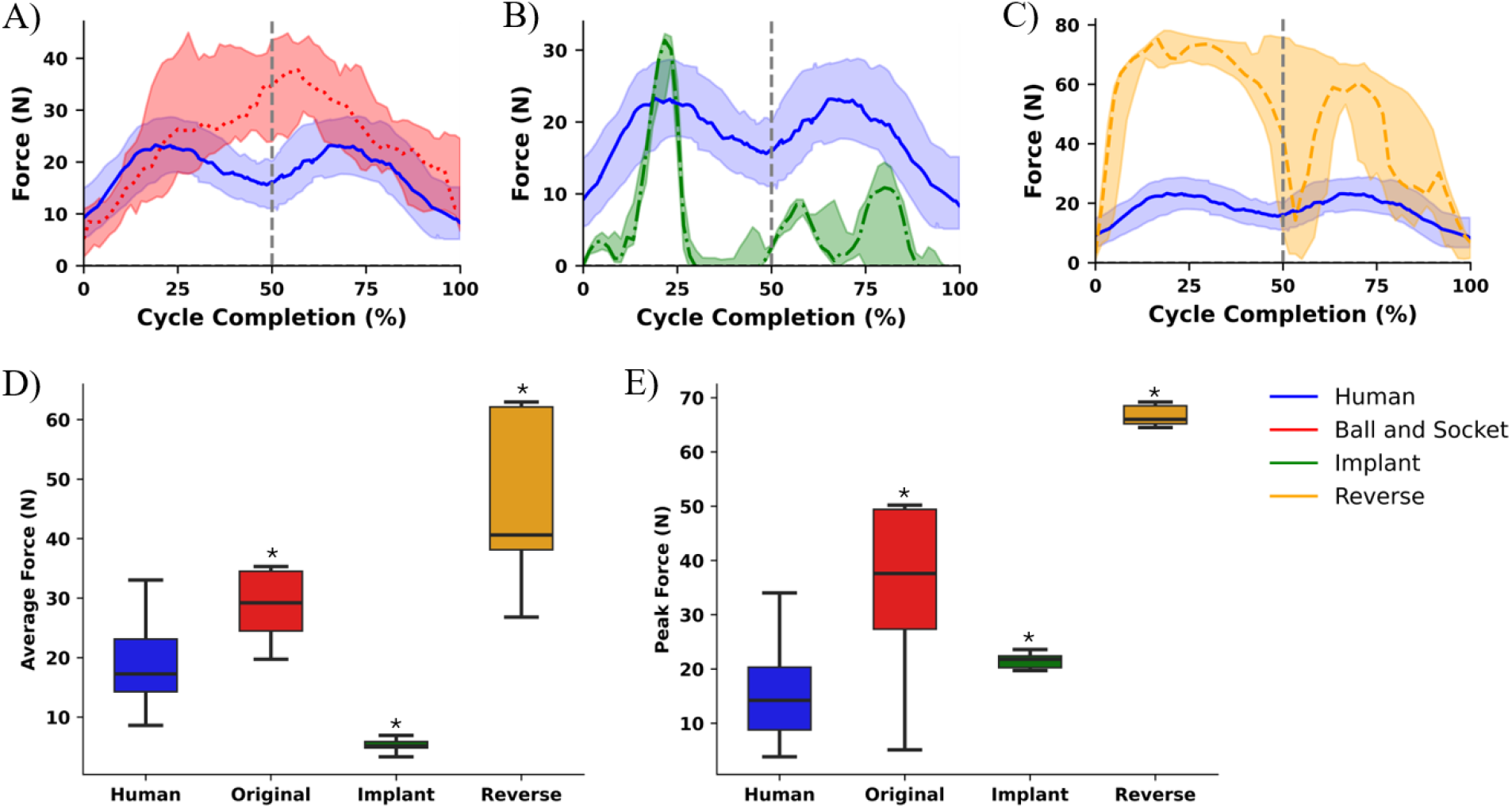
Supraspinatus force profile and values for abduction to 90° in scapular plane, measured in Newtons (N). (A) Compares the original ball and socket joint configuration’s force profile to that of the human shoulder. (B) Compares the reverse shoulder joint configuration and human shoulder force profiles. (C) Compares the clinical implant joint configuration and human shoulder force profiles. The blue curve and error bands represent median human participant force and IQR, respectively. The colored error bands represent the IQR for each joint configuration. D). Average force values for the human participants and the humanoid robot for each joint configuration during task completion. E). Peak force values for human participants and the humanoid robot for each joint configuration during task completion. Results which have statistically significant (p<0.05) values from human participants have been denoted with an asterisk (*). The movement cycle length is 6 seconds.

In the 90° abduction task, human participants showed peak force values at 25% and 75% into the movement cycle (Fig. 7A-C). This pattern is also seen in the clinical implant and reverse shoulder joint configurations (Fig. 7C, B), while the original joint (Fig. 7A) reached its peak force at the cycle’s midpoint during the 90° task.

In the 90° task, the implant joint configuration showed a sharp spike in force magnitude during the first quarter cycle. Negligible force readings around the midpoint were observed, with a slight increase towards the end of the cycle, forming an M shape, (Fig. 7B). The reverse shoulder configuration consistently generated higher forces than human participants for most of the cycle, with a sudden drop in force values mid-cycle. The original ball and socket joint configuration most closely matched the force magnitude of human participants.

Table 3 provides additional details on force magnitude in the 30° and 90° abduction trials for each joint configuration compared to human participants. In the 30° trial, peak force values experienced by the original and implant joints did not significantly differ from the peak force experienced by human participants in this task. For the original and implant joints, the humanoid robot exhibited slightly lower average force values, with differences of less than 10 N compared to human participants. However, the average force for the reverse shoulder configuration was twice as large as that of human participants. The peak force over the movement cycle for this joint was three times greater than the peak force for human participants during task completion.

**Table 3:**
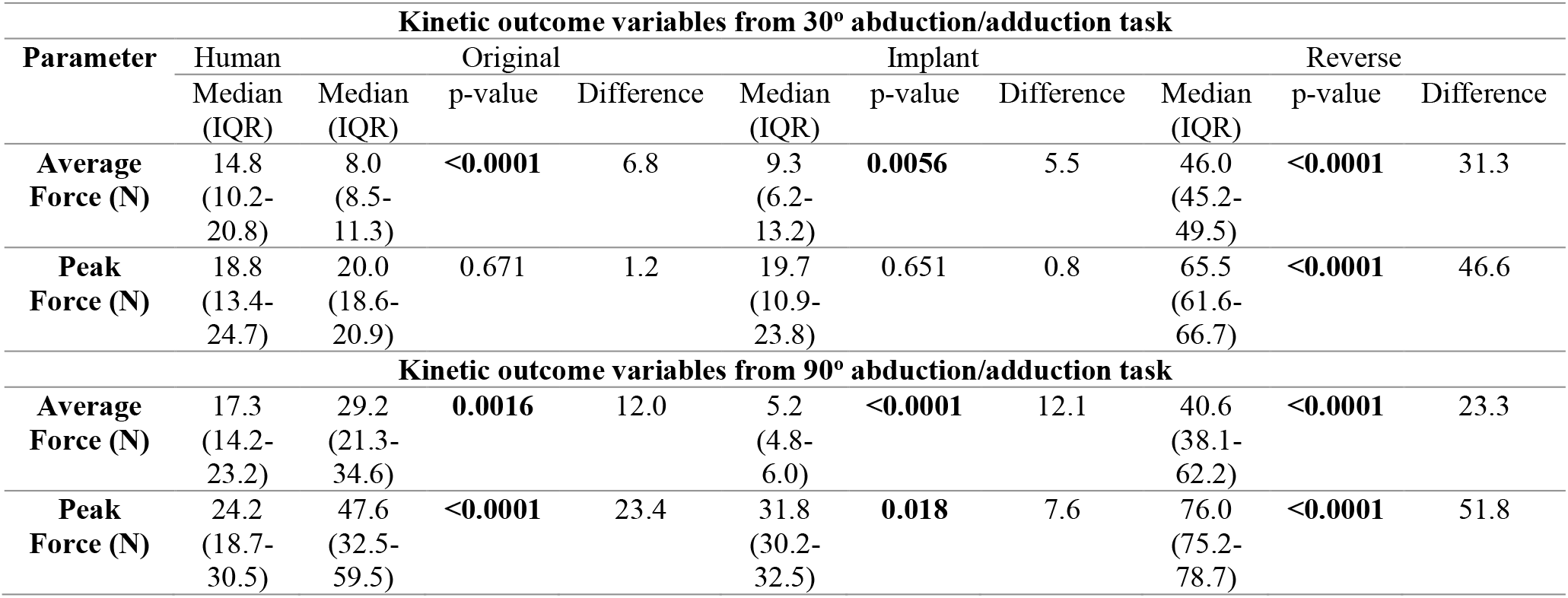
Kinetic outcome variables from 30° and 90° abduction/adduction task used to evaluate implant performance. P-values indicating that parameter of interest differs significantly from human participants (i.e., p<0.05) are bolded.

Statistically significant differences in both average and peak forces were observed between human participants and the humanoid robot in the 30° abduction trial with all three joint configurations. Peak force values for all joint configurations were higher than those of human participants during task completion. The clinical implant closely matched the peak force of human participants, with a difference of less than 10 N. The original ball and socket joint had a 23.4 N higher median peak force, while the reverse shoulder joint exceeded the median peak force of human participants by 51.8 N. The average force of human participants for the 90° abduction task was 12.1 N greater than the clinical implant joint and 12.0 N less than the original ball and socket configuration. The reverse configuration’s median average force was 23.3 N greater than the median force among human participants.

## 4. Discussion

### 4.1 Slow Movement Tasks

The humanoid robot closely replicated human kinematics during slow-paced abduction tasks up to 30° ROM. As the trial aimed to produce abduction patterns that mimic slow shoulder movement during the initial phases of supraspinatus rehabilitation, the notable difference between the mean peak acceleration values of the human and humanoid does not preclude the clinical relevance of the MSK robot to the early phase of rehabilitation. This phase of rehabilitation does not require significant accelerations, as it is focused on providing small mechanical cues to guide tendon cell alignment.

The original humanoid ball and socket joint configuration could not adequately match human abduction to 90° over the slow movement task, with statistically significant differences in peak ROM being noted. The joint’s movement over the slow 90° task was limited to an average ROM of 68.7° (SD.4.3°). It is unsurprising that the original ball and socket’s movement is constrained to around 70° ROM, given that the humanoid robot lacks key anatomic structures found in human shoulders, such as the scapula which plays an important role in overhead movements as well as protraction and retraction of the arm. For abduction between 30° and 90°, it has been estimated that the scapula contributes 20-40 % (12°-24°) of the total ROM in healthy human shoulders [46].

### 4.2 Fast Movement Tasks

Although the original ball and socket joint closely matched human movement at the slower pace, this configuration inadequately reproduced human kinematics at the higher movement speed. The clinical implant was similarly unable to replicate human performance at the higher speed. The clinical implant’s speed metric, or measure of smoothness, was found to be significantly lower than that of human participants in both trials. Hence, the implant not only completed the movements at a slower pace and to a lower peak ROM than human participants, but also moved less fluidly throughout the trajectory. This was observed qualitatively in Fig. 4B and F, with the implant quickly reaching peak ROM but having difficulty changing directions and transitioning from abduction to adduction to return to the starting point, indicating decreased joint stability.

Conversely, the reverse shoulder joint configuration closely matched human kinematics during faster-paced tasks, with minimal differences in peak ROM of less than 10° observed in both movement trials (Table 1). In the 90° abduction trial, peak ROM was the only outcome parameter found to have a statistically significant difference when compared to humans. The increased ROM and superior kinematic profile of the reverse shoulder configuration were promising findings, as this configuration is used clinically in shoulder implants to improve stability and function for patients with shoulder muscle deficiencies [47].

### 4.3 Kinetic Profiles and Joint Stability

The force magnitudes in the 30° abduction trial for the original joint and implant were similar to values estimated for human participants. No statistically significant differences in peak force and less than 10 N difference in average forces were noted. While the reverse shoulder joint was able to closely match the kinematic profile of human participants, this configuration’s mean and peak force values were consistently higher than those of human participants. The results from the 90° abduction trial, mirrored the 30° trial, with the original and implant joint configurations closely matching human force profiles, while the values in the reverse shoulder configuration experienced much higher forces in the supraspinatus position. The overall force trajectories in the 90° trial also appeared qualitatively different, with a sudden drop in force values at the supraspinatus position mid-cycle at 90°. Although the other myorobotic forces were not measured in this study, this difference in kinetic profile appears consistent with findings in cadaveric studies using reverse shoulder implants, and this finding is likely explained by the fact that at 90° ROM, other muscles such as the anterior and medial deltoid are preferentially recruited [48].

Differences in movement speeds and force profiles may be attributable to joint stability. The original joint and implant configuration were more unstable than the reverse shoulder joint, leading to greater recruitment of the myorobotic actuators in the infraspinatus, subscapularis, and teres minor positions. Movement speed was slowed by the requirement for coordination between these actuators, limiting the humanoid’s ability to mimic human kinematics for these configurations. Interestingly, this increased actuator coordination likely had the unintended effect of limiting force contributions from the supraspinatus position actuator, hence improving the kinetic mimicry of the ball and socket and implant joints.

Our results suggest that the humanoid robot used in this study can either reproduce human kinematics or provide physiologically relevant forces during task completion, but not both simultaneously. It will be important to ensure that both kinematic and kinetic profiles mimic human mechanics for applications such as tissue engineering. Adaptations will need to be made to the humanoid arm to ensure stability and an appropriate range of forces without reductions in movement speed or ROM. Future improvements to increase biomechanical and physiological relevance could include adding a scapula in the model, involving additional actuators, re-routing the tendon strings and their attachment points, and placing artificial ligaments around the joint to increase stability.

### 4.4 Limitations of study

Given that human supraspinatus muscle forces were estimated rather than directly measured and electromyography activity curves were not obtained for qualitative interpretation, the only method used to evaluate forces presented in this study was comparison with literature. Although estimated supraspinatus muscle forces reasonably matched literature values using in vitro cadaveric shoulder tissue [49, 50] and silico models [51-54], these results must be interpreted cautiously.

Our study involved a comparison with the shoulders of a young adult demographic, deviating from the conventional recruitment of middle-aged patients in randomized control trials. Our focus on a younger demographic was due to the age-related increase in rotator cuff injury prevalence [55-57]. While our study specifically examines the supraspinatus tendon, the shoulder girdle plays a multifaceted role where multiple rotator cuff muscles collaborate to ensure stability to the glenohumeral joint and facilitate movement. Human shoulders are governed by four articulating joints, numerous ligaments, and over fifteen controlling muscles [58, 59]. Human shoulders, thus, differ anatomically from the humanoid shoulder, featuring a single joint approximating the glenohumeral joint stabilized by seven myorobotic actuators mimicking the positions of the rotator cuff muscles. Consequently, the humanoid shoulder is restricted in its range of motion, unable to replicate complex rotational actions such as reaching behind the back. The reduced number of actuators and ligamentous structures in the robot further limits the precision of its movements and load bearing capabilities, as human shoulders perform highly coordinated and complex tasks through the interplay of numerous muscles and joints, a capability that robots with fewer actuators may be unable to match. Additionally, the human shoulder’s ability to make real-time continuous adjustments through sensory feedback pathways is not present in the humanoid model, thus limiting its performance in environments requiring constant adaptation and balance. Given their robust ligamentous and muscular support, human shoulders are capable of bearing and stabilizing loads across a wide ROM. As the humanoid robot is more limited in its ligament and actuator supports, it may struggle to balance loads over the same ROMS. An extension of this exploratory study may include reviewing the kinematic and kinetic properties of a robot with additional anatomic features such as a scapula over a broader ROM.

This study was completed without the use of advanced computational tools like finite element analysis and multi-objective optimization algorithms that may allow for tendon string and myorobotic actuator positioning be optimized for maximum kinetic and kinematic performance. These tools can simulate and analyze various configurations to identify the optimal placement of actuators and tendon strings that most closely replicate the biomechanics of human muscles and tendons. Future studies may review the impact of these optimization methods on robot performance.

It is also important to note that the robotic shoulder model used in this study did not include a flexible bioreactor chamber, as proposed by Mouthuy et. al [5]. These chambers provide an environment for tissue grafts to develop while undergoing physiologically relevant stress from the robotic arm. Aside from allowing for the analysis of the mechanical forces acting on the bioreactor chamber within the shoulder system, the inclusion of the chamber with its associated tendon within the robotic shoulder model may have impacted the shoulder’s function. The robotic shoulder’s precise movements and controlled forces would provide robust mechanical stress to the tissue graft. Hence, the tissue would be conditioned under dynamic conditions that mimic human kinematic and kinetic profiles.

Exposure to these mechanical stressors may lead to a dynamic tissue growth response whereby the tendon tissue develops certain functional characteristics such as elasticity and strength in a manner that does not only optimize its function within the robotic system, but that also more closely mimics the tendon histology observed in vivo [5].

### 4.5 Implications of Findings

This is a proof-of-concept study which has compared the biomechanical profile of a humanoid robot to human participants while completing abduction/adduction tasks in real-time. To date, no bioreactor systems presented in peer-reviewed literature have attempted to provide mechanical stimulation through movement patterns that consider the anatomic location of the tendon tissue being cultured [60-82]. Furthermore, many recent studies have presented bioreactor systems that provide less than 5 N of force during peak loading, which does not sufficiently match the estimated forces expected in humans [60, 62, 73, 76]. With actuators positioned to simulate rotator cuff muscles and tendons, the humanoid shoulder demonstrated the potential to replicate controlled abduction tasks at a low ROM. This is particularly relevant to shoulder tendon tissue engineering, given that the supraspinatus tendon, which is most commonly injured, is primarily responsible for these movement tasks [34]. The kinetic modelling presented in this study is intended to provide a starting point for comparison of human forces with those of the humanoid robot. Though this study demonstrated the ability of the humanoid robot to replicate controlled abduction movement tasks up to a ROM of 30°, more sophisticated models that are required prior to the clinical application of robotic tissue bioreactors [83-86]. To increase ROM and enhance the performance of this system, future prototypes of the robotic shoulder should incorporate a scapular component. This is already being developed in other MSK robots [87].

In addition to tissue engineering applications, an improved version of this MSK robot could provide more mechanically appropriate loading conditions to complement current methods of implant testing. Though this study was able to effectively replicate slow controlled abduction movements up to ROM of 30°, the addition of a scapular component and other anatomic features may allow the ability of impacts to be tested over a robust ROM that more closely mimics that of humans. Traditional biomechanical evaluation of shoulder implants relies on uniaxial and biaxial loading systems that do not comprehensively replicate in vivo loading conditions [88, 89]. While shoulder joint mechanical platforms are available, they do not effectively mimic the biological environment or the muscle-tendon-bone actuation mechanism found in humans[5]. This limitation hinders their ability to provide accurate simulations of shoulder mechanics and responses to physical stresses, thereby reducing their applicability in replicating true human shoulder function.

## 5. Conclusion

The kinematic results from this study demonstrate that the humanoid robot can effectively replicate slow, controlled abduction movements up to a ROM of 30 degrees, the ROM over which the supraspinatus is most actively engaged. This finding is particularly relevant in tissue tendon engineering, where accurate mechanical loading is crucial for the development and maturation of tissue grafts. However, to realize the full potential of humanoid robots for these applications, further work is necessary to refine the estimation of forces, future research should prioritize enhancing the robot’s actuation system to achieve precise and adaptable force control, allowing the robot to operate across a wider range of movement speeds and ROM. Such improvements will allow the robot to provide a robust mechanical environment that more closely mimics that of humans to support the development of functional tissue grafts.

## Acknowledgments

We gratefully acknowledge Devanthro GmbH, for their technical support in developing and modifying the musculoskeletal humanoid robot used in this study.

## Author contributions

I.S.: Adaptations of robotic arm, experimental design, data collection, write up; A.S.: data analysis, write up; J.S.: Experimental design, motion study protocol and coding, write up and technical expertise; A.C.: clinical expertise, write up; P.M.: Adaptations of robotic arm, experimental design, coordination of study, robotic expertise, write up

